# Effects of Malaise trap spacing on species richness and composition of terrestrial arthropod bulk samples

**DOI:** 10.1101/2020.09.30.321430

**Authors:** D Steinke, TWA Braukmann, L Manerus, A Woodhouse, V Elbrecht

**Affiliations:** Centre for Biodiversity Genomics, University of Guelph, 50 Stone Road East, Guelph, Ontario, N1G 2W1, Canada; Department of Integrative Biology, University of Guelph, 50 Stone Road East, Guelph, Ontario, N1G 2W1, Canada; Optimist Club of Kitchener-Waterloo; Centre for Biodiversity Monitoring, Zoological Research Museum Alexander Koenig, Bonn, Germany

**Keywords:** Metabarcoding, insects, biodiversity, biomonitoring, experimental design

## Abstract

The Malaise trap has gained popularity for assessing diverse terrestrial arthropod communities because it collects large samples with modest effort. A number of factors that influence collection efficiency, placement being one of them. For instance, when designing larger biotic surveys using arrays of Malaise traps we need to know the optimal distance between individual traps that maximises observable species richness and community composition. We examined the influence of spacing between Malaise traps by metabarcoding samples from two field experiments at a site in Waterloo, Ontario, Canada. For one experiment, we used two trap pairs deployed at weekly increasing distance (3m increments from 3 to 27 m). The second experiment involved a total of 10 traps set up in a row at 3m distance intervals for three consecutive weeks.

Results show that community similarity of samples decreases over distance between traps. The amount of species shared between trap pairs shows drops considerably at about 15m trap-to-trap distance. This change can be observed across all major taxonomic groups and for two different habitat types (grassland and forest). Large numbers of OTUs found only once within samples cause rather large dissimilarity between distance pairs even at close proximity. This could be caused by a large number of transient species from adjacent habitat which arrive at the trap through passive transport, as well as capture of rare taxa, which end up in different traps by chance.

## Introduction

> *During my extensive travels I have repeatedly found that insects happened to enter my tent, and that they always accumulated at the ceiling-corners in vain efforts to escape at that place without paying any attention to the open tent-door. On one occasion one of the upper tent-corners happened to have a small hole torn in the fabric, and through this hole all the insects pressed their way and escaped. Later on the idea occurred to me, that, if insects could enter a tent and not find their, way out, and always persistently tried to reach the ceiling, a trap, made as invisible as possible and put up at a place where insects are wont to patrol back and forth, might catch them much better than any tent and perhaps better than a man with a net…*
>
> — *Rene Malaise 1937*

The inclusion of terrestrial invertebrates in biodiversity inventories and surveys has increased substantially over the past years (Dopheide et al. 2019, Drake et al. 2007) but sampling efficiency remains a key consideration when designing larger censuses (Telfer et al. 2015, Timms et al. 2012). Although no single sampling can be used to survey all taxa at a given site, the Malaise trap (Malaise 1937) has gained popularity for assessing terrestrial arthropod communities (Karlsson et al. 2005) because it collects large and diverse samples with fairly little effort. Malaise’s (1937) invention is a tent-like flight-interception trap made from fine mesh netting with a central screen suspended below a sloping ridge-roof that leads to a collecting bottle at the upper end. Flying insects that hit the screen, subsequently fly or walk along this roof to the bottle which is usually filled with >90% ethanol as preservative. Traps are usually deployed in a way that the central mesh intercepts the flying path of insects. There are a number of different designs available although the most commonly used traps are so called Townes-Style traps (Townes 1972) and derived versions of it (e.g. ez-Malaise traps). The trap is particularly well suited for inventory because it catches a wide variety of flying insects and some ground active insects that climb up the trap fabric. Malaise trapping is easy, requires modest labour and as such represents one of the best mass-collecting methods available for terrestrial invertebrates (deWaard et al 2018).

Initially Malaise traps were considered of limited use in conservation evaluation and bio-surveillance because of the huge size of their catch (Drake et al. 2007) which made it difficult to characterize the community using traditional morphology-based methods (Cook et al. 2010). Consequently, larger surveys used total biomass rather than detailed identification of specimens. In fact one of the recent reports on the dramatic decline of terrestrial arthropod abundance was the result of a long-term study using Malaise traps and catch biomass (Hallmann et al. 2017). The recent advent of DNA barcoding (Hebert et al. 2003) and metabarcoding (Taberlet et al. 2012) opened the door to more comprehensive estimates of species richness and community composition (Braukmann et al. 2019, Yu et al. 2012, Steinke et al. 2020) and Malaise traps are poised to become a ubiquitous tool for biodiversity surveys (Geiger et al. 2016). It seems that large scale or global high-resolution monitoring networks are within our reach (Hobern & Hebert 2019) but there are still a number of challenges in relation to the small scale variability of many terrestrial habitats.

There are various factors that influence the efficiency of Malaise traps. Temperature, precipitation, and wind are considered important as largest catches generally occur on hot, dry, and still days (Matthews & Matthews 1971). It has also been noted (Townes 1962) that insects often fly closer to the ground in spring because of the warmer air there, thereby increasing the number of individuals caught during this season. As a Malaise trap samples only those arthropods that happen to fly through a relatively small area, trap placement becomes an important consideration. Height of surrounding vegetation and location in shade or sun can alter trap performance and efficiency (Matthews & Matthews 1971, Ssymank et al. 2018). Another relevant but not systematically studied variable is the distance between traps in a sampling area. This is a particular important consideration when designing larger biotic surveys using arrays of traps. For instance, it is not known how many traps at what distance are needed to maximise observable species richness and community composition for a given location.

The main objective of this study was to examine the effects of spacing between traps on species richness and composition of Malaise trap samples. Bulk samples from two field experiments at a site in Waterloo, Ontario, Canada were assessed using metabarcoding to determine if (1) there is a critical distance between traps at which species overlap drops significantly and if (2) structural composition of habitats has an influence on such a distance.

## Materials & Methods

### Site and sampling

Arthropod bulk samples were collected using ez-Malaise traps (Bugdorm, Taiwan). Traps for the first experiments (Figure 1a) were deployed in both a grassland and a forested pond area near Waterloo, Ontario, Canada. Traps for experiment 2 (Figure 1b,c) were positioned only in the grassland area. For the first experiment we used two trap pairs that were deployed next to each other (3m distance between both collecting bottles) respectively. Each week trap spacing for each pair was increased by three meters to a maximum distance of 27m (Figure 1a). Samples were collected every week before moving one trap further away. Each time we cleaned trap heads (collecting area with bottles) using bleach and ethanol to minimize cross-contamination between sampling events. The second experiment involved a total of 10 traps set up in 3m distance intervals for three consecutive weeks (Figure 1b,c). Samples were also collected each week sample bottles were stored at −20 °C for further analysis.

**Figure 1:**
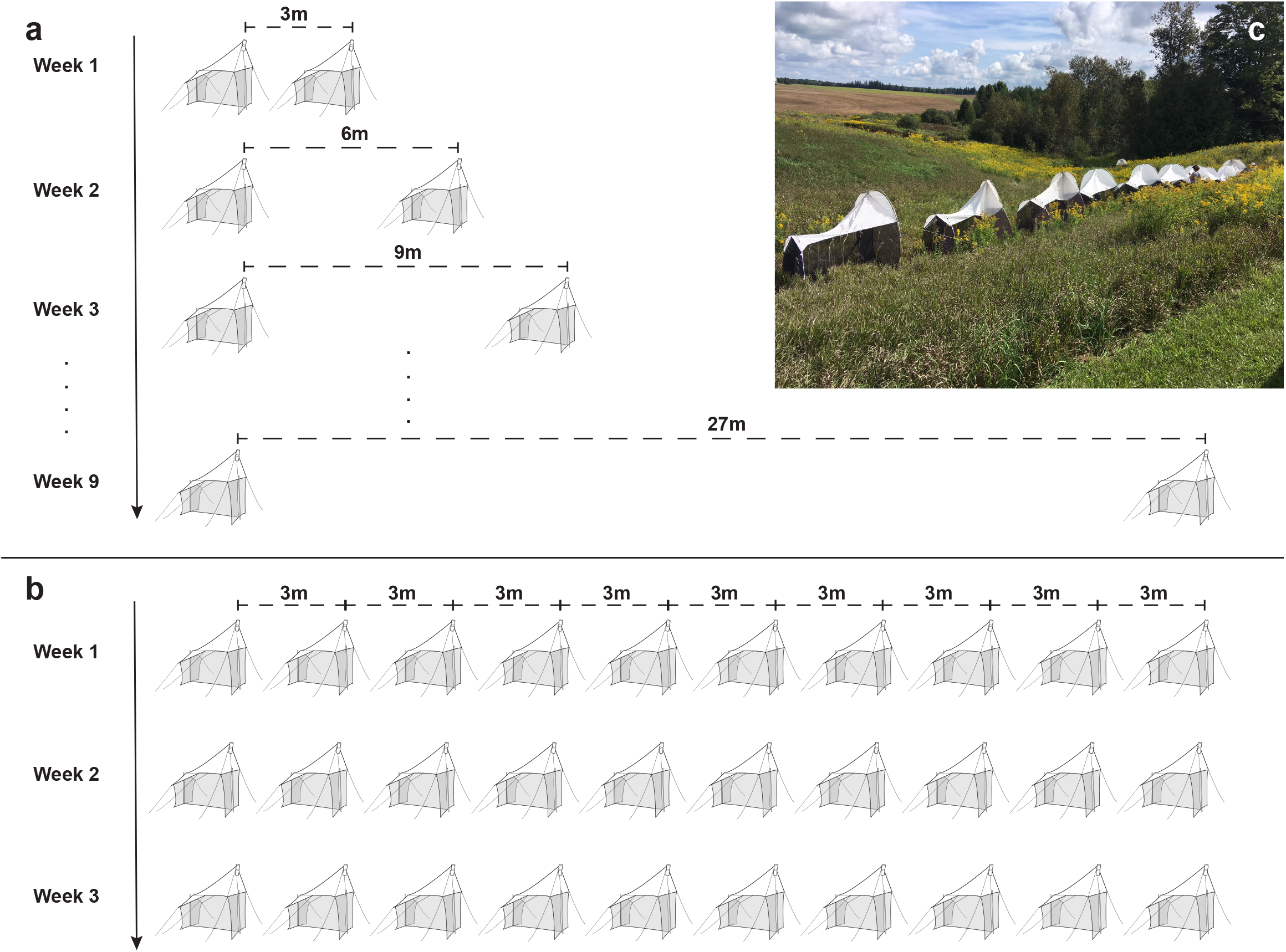
Sampling design (trap distances over time) for experiment 1 (a) and experiment 2 (b).

### Molecular analysis

All samples were dried at room temperature for three days in a disposable grinding chamber. Each sample was ground to fine powder using an IKA Tube Mill control (IKA, Breisgau, Germany) at 25,000 rpm for 2 × 3 min. DNA was extracted from approximately 20 mg of ground tissue using the DNeasy Blood & Tissue kit (Qiagen, Venlo, Netherlands) following manufacturer protocols.

Metabarcoding was carried out using a two-step fusion primer PCR protocol (Elbrecht & Steinke 2019). During the first PCR step, a 421 bp region of the Cytochrome c oxidase subunit I (COI) was amplified using the BF2 + BR2 primer set (Elbrecht & Leese 2017, Elbrecht et al. 2019). PCR reactions were carried out in a 25 μL reaction volume, with 0.5 μL DNA, 0.2 μM of each primer, 12.5 μL PCR Multiplex Plus buffer (Qiagen, Hilden, Germany). The PCR was carried out in a Veriti thermocycler (Thermo Fisher Scientific, MA, USA) using the following cycling conditions: initial denaturation at 95 °C for 5 min; 25 cycles of: 30 sec at 95 °C, 30 sec at 50 °C and 50 sec at 72 °C; and a final extension of 5 min at 72 °C. PCR success was checked on a 1% agarose gel. One μL of PCR product was used as template for the second PCR, where Illumina sequencing adapters were added using individually tagged fusion primers (Elbrecht & Steinke 2019). Tagging combinations are available in Table S1. We mainly used the same thermocycler conditions as in the first PCR but the reaction volume was increased to 35 μL, the cycle number reduced to 20 and extension time increased to 2 minutes per cycle. PCR success was again checked on a 1% Agarose gel. PCR products were purified and normalized using SequalPrep Normalization Plates (Thermo Fisher Scientific, MA, USA, Harris et al., 2010) according to manufacturer protocols. Ten μL of each normalised sample were pooled, and the final library cleaned using left sided size selection with 0.76x SPRIselect (Beckman Coulter, CA, USA. Sequencing was carried out by the Advances Analysis Facility at the University of Guelph using the 600 cycle Illumina MiSeq Reagent Kit v3 and 5% PhiX spike in. The read length of read one was increased to 316 bp, while keeping read 2 to 300 bp. As we only used inline barcodes for sample tagging, both Illumina indexing read steps were skipped.

### Data processing

Initial quality control of raw sequence data was done using FastQC v0.11.8. Subsequently, sequence data were processed using the JAMP pipeline v0.69 (github.com/VascoElbrecht/JAMP) starting with demultiplexing, followed by paired-end merging using Usearch v11.0.667 with fastq_pctid=75 (Edgar 2010). Primer sequences were trimmed from each sequence using Cutadapt v1.18 with default settings (Martin 2011), retaining only sequences where primers were successfully trimmed at both ends. Cutadapt was also used to remove sequences shorter than 411 bp and longer than 431 bp. Sequences with poor quality were removed using an expected error value of 1 (Edgar & Flyvbjerg 2015) as implemented in Usearch. Filtered reads of each sample were dereplicated and singletons removed, before pooling all reads for OTU clustering with Usearch cluster_otus at a 97% similarity threshold. Duplicated reads from each sample including singletons were mapped back against generated OTUs using Usearch usearch_global, to generate a OTU table. The maximum read count for each OTU across all 12 negative controls was multiplied by two, and subtracted from corresponding OTU read counts in all samples. Taxonomy was assigned by using OTUs as queries for the BOLD reference database (www.boldsystems.org Ratnasingham & Hebert 2007) utilizing the JAMP Bold_web_hack script with default settings. Only OTUs with a minimum match of 98% were retained for further analysis. For most analysis carried out in R v3.5.1, relative read counts were used, and only reads above 0.01% abundance were considered.

### Statistical analysis

OTU tables (Table S2) were used to calculate both the number of OTUs shared as well as the Bray-Curtis dissimilarity between all trap pairs for both experiments. In order to determine OTU sampling effort we calculated accumulation curves for both experiments using the function *specaccum* and extrapolated species richness for each week using *specpool*, both part of the vegan package (Oksanen et al. 2018). Pairwise OTU overlap among trap distance pairs was evaluated using the nonparametric multiple comparison function implemented in the R package dunn.test 1.2.4 (Dinno 2016) which is equivalent to the Kruskall-Wallis test.

## Results

We were able to extract high quality DNA from most samples, and obtained strong bands for all 74 samples after the second PCR step (Suppl Figure). Illumina sequencing generated 13,910,614 reads (partial run shared with other projects), with the raw data being available on NCBI SRA with the accession number SRP200574. About 27% of the reads were filtered during data processing, leaving an average of about 137,181 sequences per sample. In total, 10,151,381 post-filtering reads could be used for clustering with Usearch.

Our analysis shows a total of 2,315 OTUs for the grassland site and 2,804 OTUs for the forest pond site in experiment 1 (Figure 2a). On average about half of those (49%) were only detected once over the entire experiment. The Chao 1 (Magurran, 2003) species estimates for the total number of OTUs possible with complete sampling were 3,847±119 and 4,550±129 respectively. Both sites had a total of 860 OTUs in common. Bray-Curtis dissimilarity between samples of distance pairs was generally high (>0.67) for both sites and all distances, however dissimilarity at both sites increased significantly (Kruskal-Wallis and Dunn’s posthoc p < 0.0001) at 18m distance (Figure 2b). Overall, the proportion of OTUs shared between trap pairs ranged from 26-27%.

**Figure 2:**
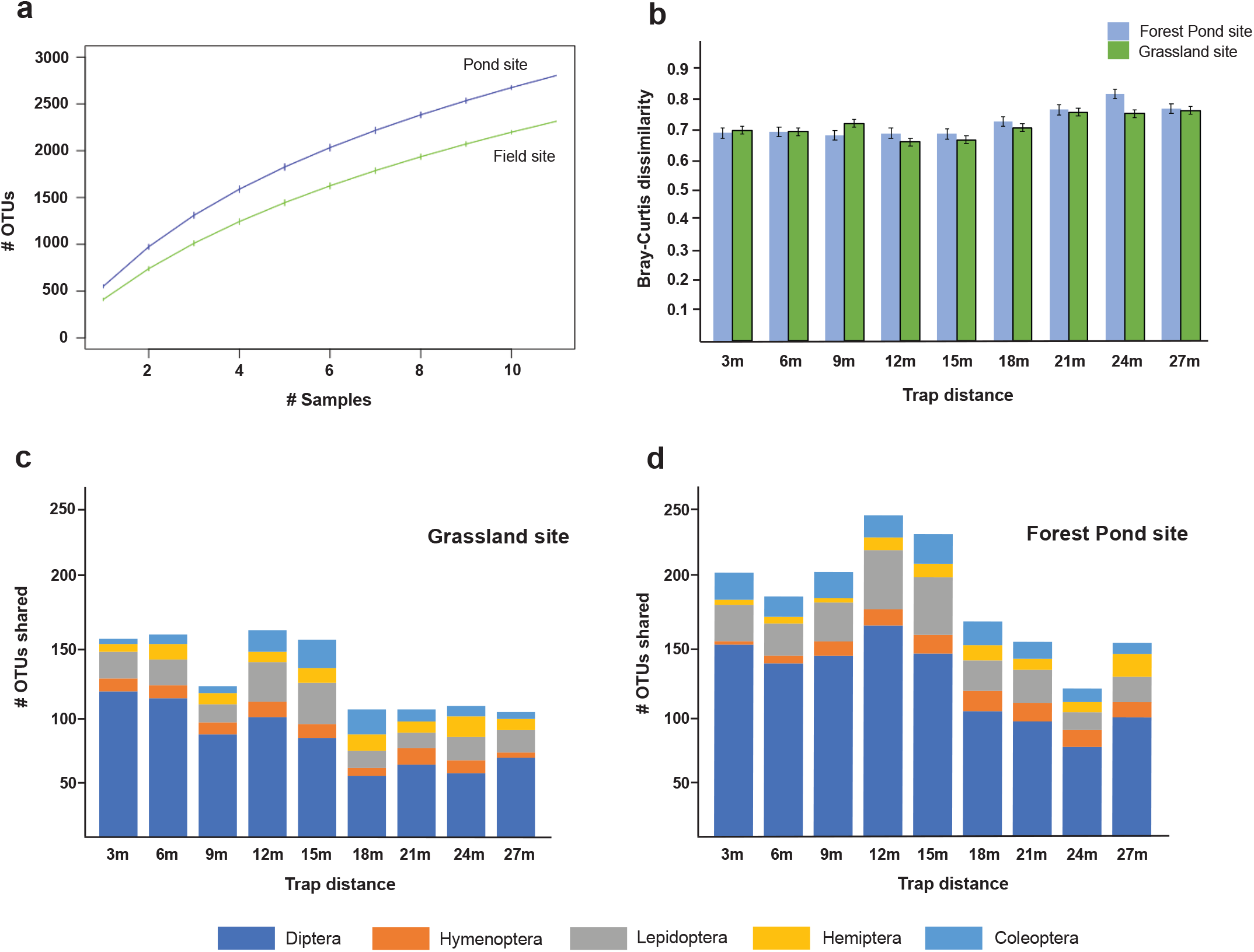
Results for experiment 1. a) OTU accumulation curves for both sites by sample, b) Histogram of Bray-Curtis dissimilarities between samples of distance pairs for both sites. Number of shared OTUs per trap distance pair for the top five arthropod orders at grassland (c) and forest pond (d) site.

The total OTU count for the grassland site comprised 21 orders with six orders (Coleoptera, Diptera, Hemiptera, Hymenoptera, Lepidoptera, Orthoptera) representing 97% of all specimens (Table S1). We found 20 orders at the forest pond site with five orders (Coleoptera, Diptera, Hemiptera, Hymenoptera, Lepidoptera) representing 96% of all specimens (Table S3). About 1/3 of OTUs was shared between both sites. We observed a distinct drop in the number of species shared between traps at a distance of 15m for both the grassland (Figure 2c) and the forest pond site (Figure 2d).

For experiment 2 we found totals of 1,017, 662, and 738 OTUs for weeks 1-3 (Figure 3a, Figure S1). The total number of OTUs found over the three weeks was 1,610. Chao estimates for the expected amount of total OTUs were 2,007+117, 1,211±80, 1,479±102 for weeks one to three, respectively. Dissimilarity values for experiment 2 were generally higher with weekly averages ranging from 0.75 (week 2) to 0.92 (week 3) (Figure 3b). The values gradually increased with increasing distance between two traps. In addition more than half of the OTUs obtained in in three weeks of experiment 2 (811) were only detected in a single trap following a common hollow curve species abundance pattern (Figure S2).

**Figure 3:**
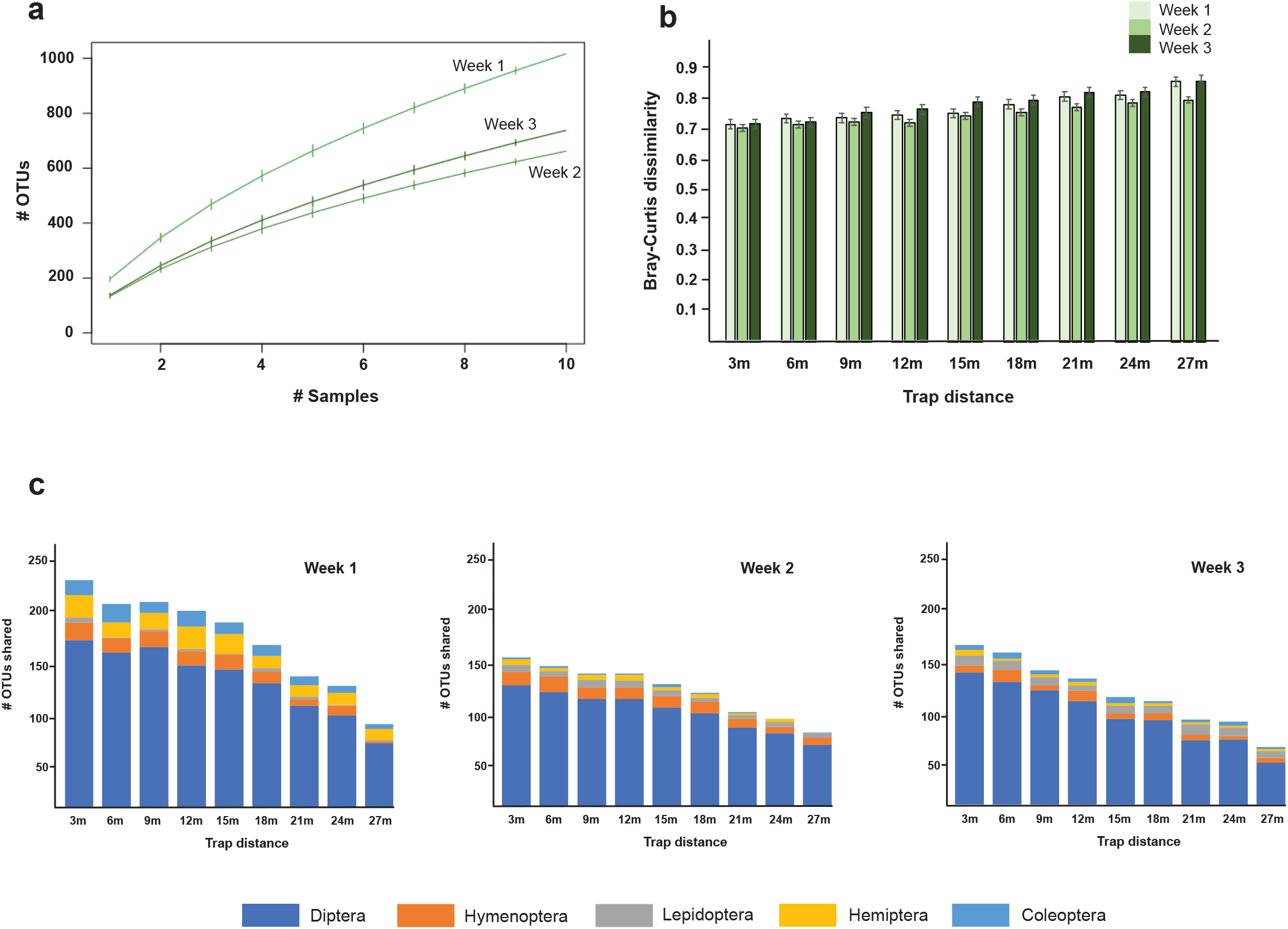
Results for experiment 2. a) OTU accumulation curves for each sampling week by trap b) Histogram of Bray-Curtis dissimilarities between samples of distance pairs for all weeks, c) number of shared OTUs by trap distance for the top five arthropod orders for each week.

OTUs found during experiment 2 comprised 15-18 orders with the five orders Coleoptera, Diptera, Hemiptera, Hymenoptera, and Lepidoptera representing between 85-95% of all specimens (Table S3). In contrast to experiment 1, each week we observed a constant decline of the number of species shared between traps with increased distance between them (Figure 3c).

## Discussion

Malaise traps as sampling method for terrestrial arthropod communities represent a rather efficient and economical means for obtaining comprehensive samples with minimal effort (Karlsson et al. 2005). They can be operated continuously in any weather with only occasional attendance and deliver large sample sizes. In conjunction with modern DNA-based methods to assign taxonomy (e.g. metabarcoding) they probably represent the best mass-collecting method available for terrestrial arthropods and are well suited for large scale biotic surveys using arrays of traps (Yu et al 2012, deWaard et al. 2018, Steinke et al 2020). However, so far it was not known how many traps at what distance are needed to maximise observable species richness and community composition for a given location (Noyes 1989). Various strategies have been applied but trap spacing varied considerably (Fraser et al. 2008, Santos et al. 2014). Results from our first experiment (Figure 1a, 2) suggest that deploying traps at about 15m distance from each other would significantly increase overall species richness and reduce overlap between traps. This is true for all major taxonomic groups collected (Figure 2 c, d). Interestingly, overall general habitat structure seems to have no effect on the distance observed as both, the grassland and the forest pond sites exhibit the same cut-off value. On the other hand, experiment 2 does not show such a clear drop in overlap between adjacent traps (Figure 3 c, d, e). For each week we were able to observe a more gradual decline in the number of OTUs overall and per taxonomic group respectively. This could be the result of the experimental set up we chose. A row of ten traps represents a continuous structure along which some animals have the ability to move before being caught. Along the row of traps the amount of observed OTUs varies which is likely the result of variation in microhabitat structure (Figure S1). The grassland chosen for the experiment was not entirely uniform and characterized by sporadic patches of golden rod (*Solidago canadensis*).

The large dissimilarity values observed in both our experiments (Figure 2b, 3b) are influenced by a large proportion of singleton OTUs. We are confident that these are mostly true specimens rather than OTUs derived from sequencing or PCR errors, because we removed OTUs that did not match the BOLD database to at least 98%. Large numbers of OTUs found only once over a sampling period or between traps have been observed several times in other studies using Malaise traps (e.g. Geiger et al. 2016, deWaard et al. 2018, Steinke et al. 2020). This phenomenon has been discussed as an indicator for the presence of transient species (D’Souza & Hebert 2018, Steinke et al. 2020). Transient species have been defined as species that show up only occasionally as a result of dispersal from adjacent habitat (Snell Taylor et al. 2018). Specifically, many smaller species caught are not necessarily living in the sampled habitat but are rather passively transported there e.g. by wind. Additionally, sampling might be stochastic when it comes to rare or low abundant taxa.

Malaise trapping with only a few traps at a single site over a short timescale always provides an incomplete species list. That is no different for our study which suggests that additional trapping efforts by increasing the number of traps or by enlarging the trapping surface (e.g. Gressitt & Gressitt 1962) are needed to approach asymptotic species richness at both experimental sites. The trap results for experiment 1 suggest that it needs a 1.6-fold increase of the full sampling effort for a complete inventory (over the entire 10 weeks of the experiment) based on Chao-1. For experiment 2 sampling efforts would need to be doubled to obtain maximum species richness for the site in any given week. This could perhaps be accomplished by deploying a second row of ten traps at 15 m distance following the findings of experiment 1. The alternative would be to increase the sampling duration (Fraser et al. 2008) or the sampling surface of the traps (Gressitt & Gressitt 1962).

In conclusion, our results suggest the following recommendations for sampling and monitoring terrestrial invertebrate communities with Malaise traps: (a) within a temperate and uniform habitat a number of traps equally spaced at >15m will sample more of the local diversity while at the same time reduce the extend of repetitive sampling, (b) longer trapping duration can help to reach asymptotic species richness and lead to more complete species lists, and (c) future work should include research on the origin and the role of singletons. Are they in fact transient species passively dispersed towards the trap or low abundant resident core species that are not efficiently detected?

## Acknowledgements

This study was supported by funding through the Canada First Research Excellence Fund. It represents a contribution to the University of Guelph Food From Thought research program. We thank the Optimist Club of Kitchener-Waterloo for access to their site at Camp Heidelberg. We are grateful to peer reviewers and editors who provided comments that improved the manuscript.

## Author contributions

DS developed the concept, DS and VE designed the experiments and carried out the sampling, VE and LM did the laboratory work, DS and VE analysed the data, DS wrote the paper, VE, AW, and TB revised the paper and provided input throughout the study.

---

**Figure S1:**
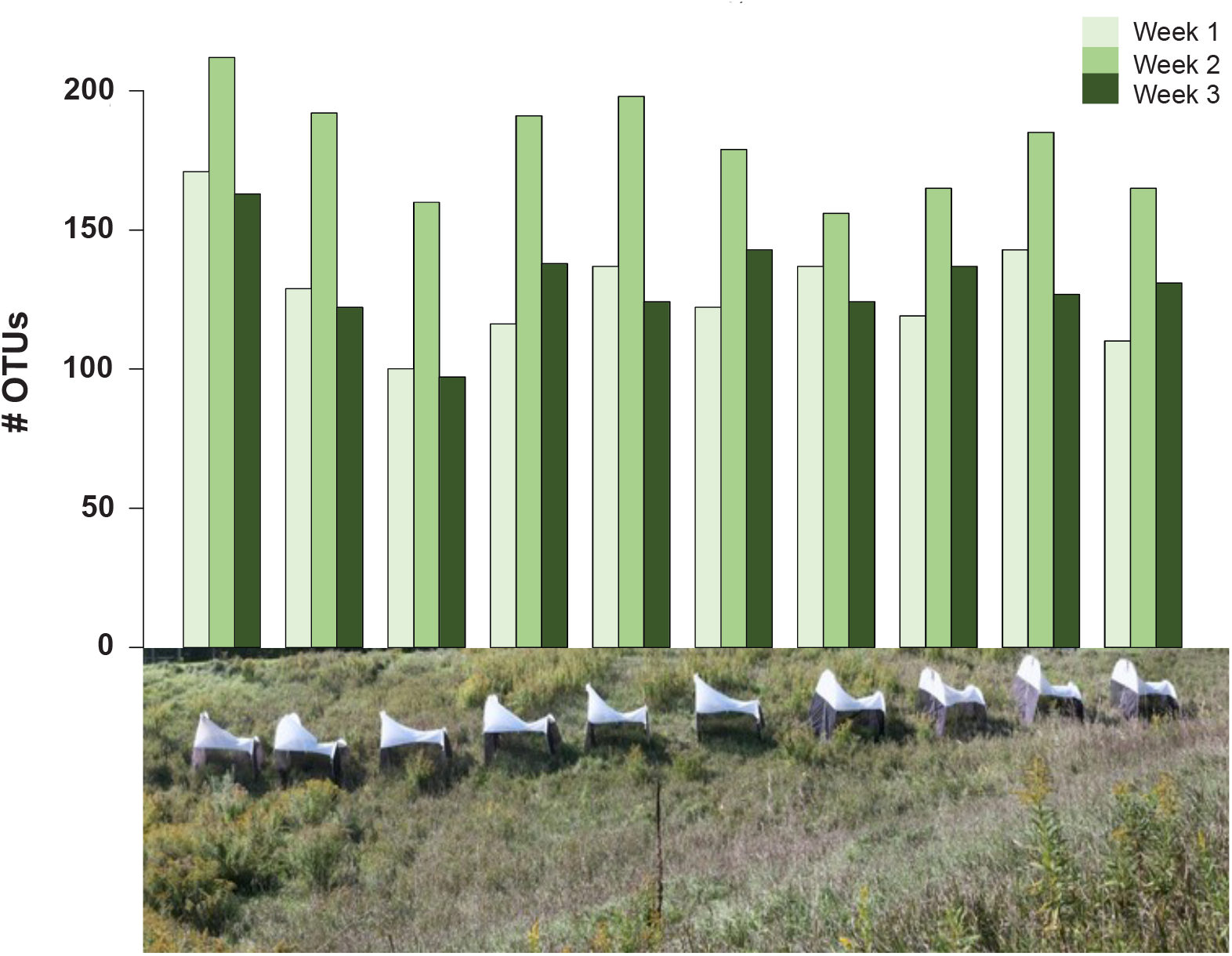
Histogram of OTU richness per trap (corresponding trap shown in photograph below) per week of experiment 2.

**Figure S2:**
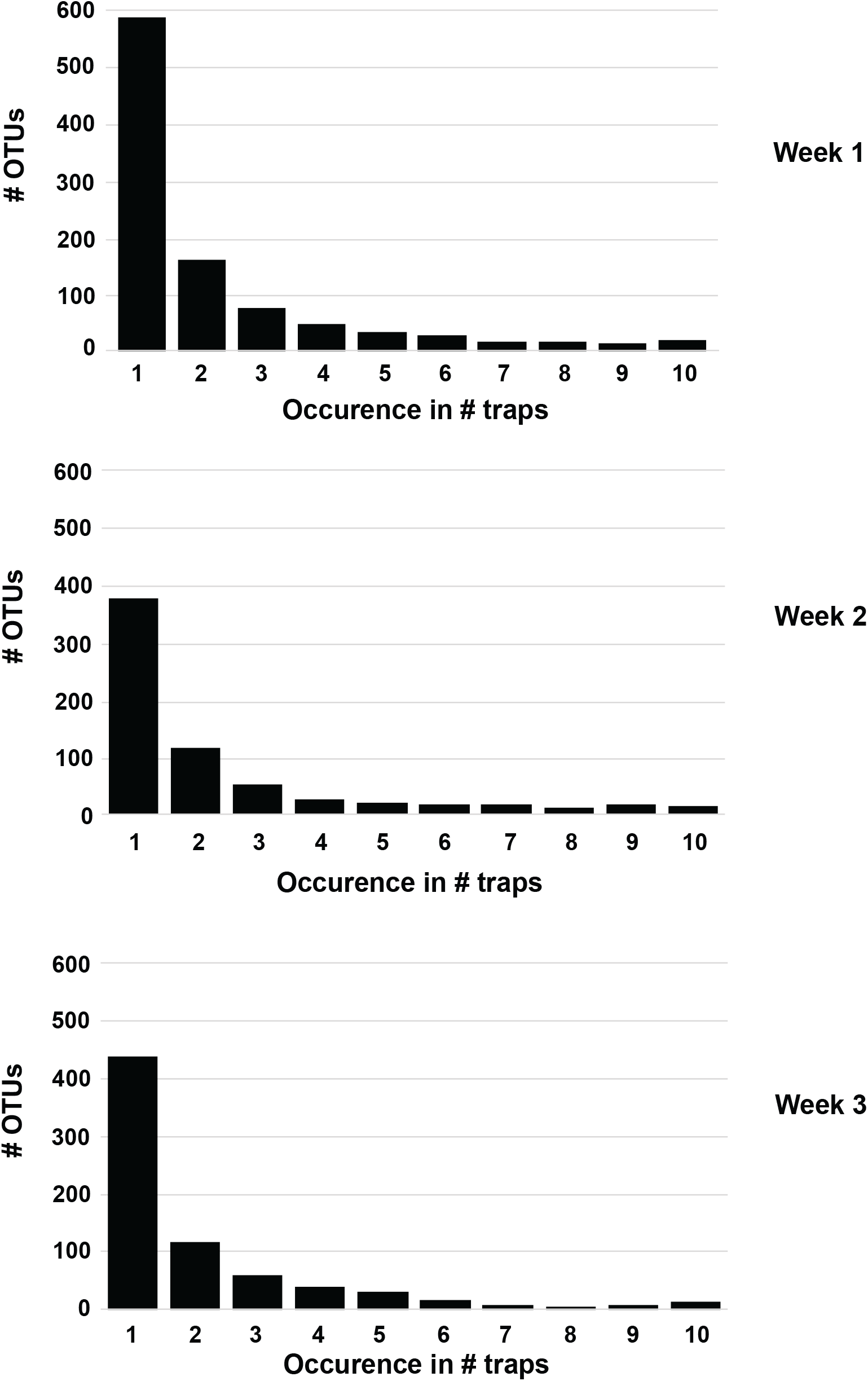
Number of OTU occurrences in traps of experiment 2.

**Table S1:** Tagging layout

*TableS1_Distance Layout.xls*

**Table S2:** OTU table (FC, FM, PC, PM – Experiment 1; A, B, F – Experiment 2; C – Controls)

TableS2_OTU Table.xlsx

**Table S3:** Taxonomic breakdown by order for both experiments and sites

TableS3_taxonomic breakdown.xlsx

## References

Beng KC, Tomlinson KW, Shen XH, Surget-Groba Y, Hughes AC, Corlett RT, Slik WF (2016). The utility of DNA metabarcoding for studying the response of arthropod diversity and composition to land-use change in the tropics. Scientific Reports, 1–13.

Braukmann TWA, Prosser SJR, Ivanova NV, Elbrecht V, Steinke D, Ratnasingham R, deWaard JR, Sones JE, Zakharov EV, Hebert PDN (2019). Metabarcoding a Diverse Arthropod Mock Community. Molecular Ecology Resources 19: 711–727.

Bush A, Sollmann R, Wilting A, Bohmann K, Cole B, Balzter H, Martius C, Zlinszky A, Calvignac-Spencer S, Cobbold CA, Dawson TP, Emerson BC, Ferrirer S, Gilbert MTP, Herold M, Jones L, Leendertz FH, Matthews L, Millington JDA, Olson JR, Ovaskainen O, Raffaelli D, Reeve R, Rödel M-O, Rodgers TW, Snape S, Visseren-Hamakers I, Vogler AP, White PCL, Wooster MJ, Yu DW (2017). Connecting Earth observation to high-throughput biodiversity data. Nature Ecology & Evolution 1: 0176.

Cook LG, Edwards RD, Crisp MD, Hardy NB (2010). Need morphology always be required for new species descriptions? Invertebrate Systematics 24: 322–326.

deWaard JR, Levesque-Beaudin V, deWaard SL, Ivanova NV, McKeown JTA, Miskie R, Naik S, Perez KHJ, Ratnasingham S, Sobel CN, Sones JE, Steinke C, Telfer AC, Young A, Young MR, Zakharov EV, Hebert PDN (2018). Expedited assessment of terrestrial arthropod diversity by coupling Malaise traps with DNA barcoding. Genome 62: 85–95.

Dinno A (2016). dunn.test: Dunn’s Test of Mulitple Comparisons Using Rank Sums. R package version 1.3.2.. http://CRAN.R-project.org/package=dunn.test.

Dopheide A, Tooman LK, Grosser S, Agabiti B, Rhode B, Xie D, Stevens MI, Nelson N, Buckley, TR, Drummond AJ, Newcomb RD (2019). Estimating the biodiversity of terrestrial invertebrates on a forested island using DNA barcodes and metabarcoding data. Ecological Applications 29: e01877.

Drake CM, Lott DA, Alexander KNA, Webb J (2007). Surveying terrestrial and freshwater invertebrates for conservation evaluation. Natural England Research Report NERR005, Sheffield, UK.

Edgar RC (2010). Search and clustering orders of magnitude faster than BLAST. Bioinformatics 26: 2460–2461.

Edgar RC, Flyvbjerg H (2015). Error filtering, pair assembly and error correction for next-generation sequencing reads. Bioinformatics 31: 3476–3482.

Elbrecht V, Leese F (2017). Validation and development of COI metabarcoding primers for freshwater macroinvertebrate bioassessment. Frontiers of Environmental Science 5: 11.

Elbrecht V, Steinke D (2019). Scaling up DNA metabarcoding for freshwater macrozoobenthos monitoring. Freshwater Biology 64: 380–387.

Elbrecht V, Braukmann TWA, Ivanova NV, Prosser SWJ, Hajibabaei M, Wright M, Zakharov EV, Hebert PDN, Steinke D (2019): Validation of COI metabarcoding primers for terrestrial arthropods. PeerJ 7: e7745.

Fraser SEM, Dytham FC, Mayhew PJ (2008). The effectiveness and optimal use of Malaise traps for monitoring parasitoid wasps. Insect Conservation and Diversity 1: 22–31.

Geiger MF, Moriniere J, Hausmann A, Haszprunar G, Waegele W, Hebert PDN, Rulik B (2016). Testing the Global Malaise Trap Program – How well does the current barcode reference library identify flying insects in Germany? Biodiversity Data Journal 4: e10671.

Gressitt JL, Gressitt MK (1962). An improved Malaise trap. Pacific Insects 4(1): 87–90.

Hallmann CA, Sorg M, Jongejans E, Siepel H, Hofland N, Schwan H, Stenmans W, Müller A, Sumser H, Hörren T, Goulson D, de Kroon H (2017). More than 75 percent decline over 27 years in total flying insect biomass in protected areas. PLoS ONE 12 (10): e0185809.

Harris JK, Sahl JW, Castoe TA, Wagner BD, Pollock DD, Spear JR (2010). Comparison of normalization methods for construction of large, multiplex amplicon pools for next-generation sequencing. Applied and Environmental Microbiology 76: 3863–3868.

Hobern D, Hebert PDN (2019). BIOSCAN – Revealing eukaryote diversity, dynamics and interactions. Biodiversity Information Science and Standards 3: e37333.

Karlsson D, Pape T, Johanson KA, Liljeblad J, Ronquist, F (2005). The Swedish Malaise Trap Project, or how many species of Hymenoptera and Diptera are there in Sweden? Entomologisk Tidskrift 126: 43–53.

Magurran AE (2003). Measuring Biological Diversity. Wiley-Blackwell, Malden, Massachusetts. 264 pp.

Malaise R (1937). A new insect trap. Entomologisk tidskrift 58: 148–160.

Martin M. 2011. Cutadapt removes adapter sequences from high-throughput sequencing reads. EMBnet Journal 17(1):10–12.

Matthews RW, Matthews JR (1971). The Malaise Trap: Its Utility and Potential for Sampling Insect Populations. The Great Lakes Entomologist 4: 117–122.

Noyes, JS (1989). A study of five methods of sampling Hymenoptera (Insecta) in a tropical rainforest, with special reference to the Parasitica. Journal of Natural History 23: 285–298.

Oksanen J, Blanchet FG, Friendly M, Kindt R, Legendre P, McGlinn D, Minchin PR, O’Hara RB, Simpson GL, Solymos P, Stevens MHH, Szoecs E, Wagner H (2018). vegan: Community Ecology Package. R package version 2.5-1. https://CRAN.R-project.org/package=vegan.

R Core Team (2018). R: A language and environment for statistical computing. R Foundation for Statistical Computing, Vienna, Austria. URL https://www.R-project.org/.

Santos EFD, Noll FB, Brandao CRF (2014). Functional and Taxonomic Diversity of Stinging Wasps in Brazilian Atlantic Rainforest Areas. Neotrop Entomol. 43, 97–105.

Snell Taylor SJ, Evans BS, White EP, Hurlbert AH (2018). The prevalence and impact of transient species in ecological communities. Ecology 99(8): 1825–1835.

Steinke D, de Waard S, Sones, JE, Ivanova NV, Prosser SJR, Perez K, Braukmann TWA, Milton M, Borisenko, A, Elbrecht V, Zakharov EV, deWaard JR, Ratnasingham S, Hebert PDN (2020). Message in a bottle – Ecoregion biodiversity surveys using Malaise Traps. Submitted/preprint.

Ssymank A, Sorg M, Doczkal D, Rulik B, Merkel-Wallner G, Vischer-Leopold M (2018). Praktische Hinweise und Empfehlungen zur Anwendung von Malaisefallen für Insekten in der Biodiversitätserfassung und im Monitoring. Series Naturalis 1:1–12.

Taberlet P, Coissac E, Pompanon F, Brochmann C, Willerslev E (2012). Towards next-generation biodiversity assessment using DNA metabarcoding. Molecular Ecology 21: 2045–2050.

Telfer AC, Young MR, Quinn J, Perez K, Sobel CN, Sones JE, Levesque-Beaudin V, Derbyshire R, Fernandez-Triana J, Rougerie R, Thevanayagam A, Boskovic A, Borisenko AV, Cadel A, Brown A, Pages A, Castillo AH, Nicolai A, Mockford B, Mockford G, Bukowski B, Wilson B, Trojahn B, Lacroix CA, Brimblecombe C, Hay C, Ho C, Steinke C, Warne CP, Garrido Cortes C, Engelking D, Wright D, Lijtmaer DA, Gascoigne D, Hernandez Martich D, Morningstar D, Neumann D, Steinke D, DeBruin D, DeBruin M, Dobias D, Sears E, Richard E, Damstra E, Zakharov EV, Laberge F, Collins GE, Blagoev GA, Grainge G, Ansell G, Meredith G, Hogg I, McKeown J, Topan J, Bracey J, Guenther J, Sills-Gilligan J, Addesi J, Persi J, Layton KKS, D’Souza K, Dorji K, Grundy K, Nghidinwa K, Ronnenberg K, Min Lee K, Xie K, Lu L, Penev L, Gonzalez M, Rosati ME, Kekkonen M, Kuzmina M, Iskandar M, Mutanen M, Fatahi M, Pentinsaari M, Bauman M, Nikolova N, Ivanova NV, Jones N, Weerasuriya N, Monkhouse N, Lavinia PD, Jannetta P, Hanisch PE, McMullin RT, Ojeda Flores R, Mouttet R, Vender R, Labbee RN, Forsyth R, Lauder R, Dickson R, Kroft R, Miller SE, MacDonald S, Panthi S, Pedersen S, Sobek-Swant S, Naik S, Lipinskaya T, Eagalle T, Decaëns T, Kosuth T, Braukmann T, Woodcock T, Roslin T, Zammit T, Campbell T, Dinca V, Peneva V, Hebert PDN, deWaard JR (2015). Biodiversity inventories in high gear: DNA barcoding facilitates a rapid biotic survey of a temperate nature reserve. Biodiversity Data Journal 3: e6313.

Timms LL, Bowden JJ, Summerville KS, Buddle CM (2012). Does species-level resolution matter? Taxonomic sufficiency in terrestrial arthropod biodiversity studies. Insect Conservation and Diversity 6: 453–462.

Townes H (1962). Design for a Malaise Trap. Proceedings of the Entomological Society Washington 64: 253–262.

Townes H (1972). A light-weight Malaise trap. Entomological News 83: 239–247.

Yu DW, Ji Y, Emerson BC, Wang X, Ye C, Yang C, Ding Z (2012). Biodiversity soup: metabarcoding of arthropods for rapid biodiversity assessment and biomonitoring. Methods in Ecology and Evolution 3: 613–623.

